# Identification of *MYOM2* as a candidate gene in hypertrophic cardiomyopathy and Tetralogy of Fallot and its functional evaluation in the *Drosophila* heart

**DOI:** 10.1101/2020.08.18.255760

**Authors:** Emilie Auxerre-Plantié, Tanja Nielsen, Marcel Grunert, Olga Olejniczak, Andreas Perrot, Cemil Özcelik, Dennis Harries, Faramarz Matinmehr, Cristobal Dos Remedios, Christian Mühlfeld, Theresia Kraft, Rolf Bodmer, Georg Vogler, Silke R. Sperling

## Abstract

The causal genetic underpinnings of congenital heart diseases, which are often complex and with multigenic background, are still far from understood. Moreover, there are also predominantly monogenic heart defects, such as cardiomyopathies, with known disease genes for the majority of cases. In this study, we identified mutations in myomesin 2 (*MYOM2*) in patients with Tetralogy of Fallot (TOF), the most common cyanotic heart malformation, as well as in patients with hypertrophic cardiomyopathy (HCM), who do not exhibit any mutations in the known disease genes. MYOM2 is a major component of the myofibrillar M-band of the sarcomere and a hub gene within interactions of sarcomere genes. We show that patient-derived cardiomyocytes exhibit myofibrillar disarray and reduced passive force with increasing sarcomere lengths. Moreover, our comprehensive functional analyses in the *Drosophila* animal model reveal that the so far uncharacterized fly gene *CG14964* may be an ortholog of *MYOM2*, as well as other myosin binding proteins (henceforth named as *Drosophila <u>M</u>yomesin a<u>n</u>d <u>M</u>yosin Binding protein (dMnM)*). Its partial loss-of-function or moderate cardiac knockdown results in cardiac dilation, whereas more severely reduced function causes a constricted phenotype and an increase in sarcomere myosin protein. Moreover, compound heterozygous combinations of *CG14964* and the sarcomere gene *Mhc* (*MYH6/7*) exhibited synergistic genetic interactions. In summary, our results suggest that *MYOM2* not only plays a critical role in maintaining robust heart function but may also be a candidate gene for heart diseases such as HCM and TOF, as it is clearly involved in the development of the heart.

**SUMMARY STATEMENT:** *MYOM2* plays a critical role in establishing or maintaining robust heart function and is a candidate gene for heart diseases such as hypertrophic cardiomyopathy and Tetralogy of Fallot.

## INTRODUCTION

Malformations of the heart and their associated diseases, present at birth up to high adult age, are one of the leading causes of death worldwide. A heterogeneous disease with different anatomical variants, physiologic manifestations, and genetic underpinnings is hypertrophic cardiomyopathy (HCM), which predominantly causes left ventricular (LV) hypertrophy and outflow tract obstruction (Jones et al., 2017; Maron et al., 2006). The hypertrophy of LV and interventricular septum causes also problems in the heart’s electrical system, which result in life-threatening arrhythmias and an increased risk of sudden death (Driscoll, 2016). HCM can affect individuals of any age, although early manifestations are rare. It is typically caused by monogenic mutations mostly located in sarcomere genes, such as myosin-binding protein C 3 (*MYBPC3*; 30-40% of cases) and beta-myosin heavy chain 7 (*MYH7*; 30-50% of cases) (Vermeer et al., 2016).

Rare deleterious mutations in genes essential for the assembly of the sarcomere have been found in patients with Tetralogy of Fallot (TOF) (Grunert et al., 2014). Congenital heart disease (CHD) is the most common birth defect with an incidence of about 1% of all newborns worldwide (van der Linde et al., 2011), with TOF being the most common cyanotic form. While HCM mainly refers to left heart structures, TOF is a cardiac anomaly of a combination of four cardiac features, including ventricular septum defect, overriding aorta, right ventricular (RV) outflow tract obstruction and RV hypertrophy. The burden of a corrected TOF heart is often well tolerated during childhood but there is significant incidence particularly of symptomatic arrhythmias during the third postoperative decade and thereafter (Cuypers et al., 2014). In general, CHDs like TOF are complex disorders of multifactorial origin, comprising genetic and epigenetic causes as well as environmental factors that lead to structural defects and heart dysfunction (Rickert, 2016) Recently, we identified a multigenic background of rare deleterious mutations in several genes, which discriminate TOF cases from controls (Grunert et al., 2014). Here, mutations in sarcomere genes such as *MYH7* and *MYBPC3* (single TOF cases) as well as titin (*TTN*) and the M-band protein myomesin 2 (*MYOM2*) (multiple TOF cases) have been identified.

Myomesin proteins comprise three family members encoded by *MYOM1, MYOM2* and *MYOM3*, and localize to the M-band of the sarcomere. These three myomesin isoforms correlate with the contractile properties in different fiber types (Agarkova and Perriard, 2005; Gautel and Djinović-Carugo, 2016). *MYOM1* is expressed in all striated muscles, while *MYOM2* is expressed in the adult heart and fast fibers, and *MYOM3* is expressed only in skeletal muscle intermediate fiber types (Agarkova et al., 2003; Gautel and Djinović-Carugo, 2016; Schoenauer et al., 2005). Moreover, the expression of *MYOM2* is weaker in the embryonic human heart but similar to the skeletal muscle, with no expression in the smooth muscle (Schoenauer et al., 2010). In addition, cardiomyocytes derived from induced pluripotent stem cells of healthy individuals and TOF patients reveal that *MYOM1* and *MYOM2* are expressed during cardiac differentiation (Grunert et al., 2020). Furthermore, *Myom2* is also expressed in embryonic and adult mouse hearts (Grunert et al., 2014). Composed of tandems of fibronectin type III (FN3) and immunoglobulin type II (Ig) domains, they act as cross-linker for the neighboring thick filaments of myosin in the M-band and also interact with TTN as its C-terminal part converges to the M-band (Agarkova et al., 2003; Hu et al., 2015). Moreover, MYOM1 and MYOM2 also interact with other sarcomere proteins such as MYH7 (Obermann et al., 1997; Obermann et al., 1998), which is known to be involved in HCM (Maron and Maron, 2012) and other CHDs (Basu et al., 2014; Bettinelli et al., 2013; Postma et al., 2011). Besides sequence alterations, RNA splicing of sarcomere genes such as troponin T (*TNNT1, TNNT2*), troponin I (*TNNI1, TNNI3*) and *MYH7* were also found to be significantly altered in patients with ischemic cardiomyopathy (Kong et al., 2010) and TOF (Grunert et al., 2016). The connection between these two distinct phenotypes, a monogenetic disease of LV heart structures (HCM) and a multigenic disease of RV heart structures (TOF), seems to be primarily based on genomic but also on transcriptomic alterations in sarcomere genes. These alterations probably contribute to an impaired RV/LV function in both diseases, and cause in short- or long-term clinical outcome arrhythmias and other disorders.

In this study, we identified mutations in the sarcomere gene *MYOM2* in two independent cohorts of unrelated TOF and HCM patients. Interestingly, the HCM patients harbor no mutations in the 12 most common HCM disease genes. As the proportion of TOF patients with *MYOM2* mutations is quite high and *MYOM2* is not described for HCM so far, we further investigated *MYOM2* as candidate gene for TOF and HCM using patient-derived cardiomyocytes (CMs) and the *Drosophila* genetic model system. *Drosophila* has a high degree of cardiac gene conservation (e.g., NKX2-5, TBX20, GATA4/6 and HAND1/2) (Bodmer, 1993; Han and Olson, 2005; Qian and Bodmer, 2009; Qian et al., 2008) and similar cardiac developmental pathways (e.g., NOTCH signaling which is involved in many CHDs including TOF) (Zhou and Liu, 2014). Moreover, the functional conservation between the vertebrate and fly heart (e.g., autonomous contraction of CMs) (Klassen et al., 2017; Ocorr et al., 2014; Taghli-Lamallem et al., 2016; Wolf et al., 2006) enables the study of cardiac function of candidate genes for congenital and other heart diseases. Here we show for the first time that patient-derived CMs exhibit myofibrillar disarray and reduced passive force with increasing sarcomere lengths. Our *Drosophila* studies show that a fly counterpart of *MYOM2, CG14964*, control fly heart size in a gene dosage-dependent manner, regulates myosin levels and also genetically interacts with sarcomere myosin heavy chain, pointing towards an intricate mechanism between MYOM2 and the sarcomere in the context of HCM and TOF.

## RESULTS

### Identification of MYOM2 as novel candidate gene in HCM and TOF

As already mentioned, we have previously shown a multigenic background of TOF by characterizing a cohort of 13 clinically well-defined isolated TOF patients who carry combinations of rare deleterious mutations in genes essential for, among others, apoptosis and cell growth as well as the structure and function of the sarcomere (Grunert et al., 2014). The cohort comprises mutations in *MYH7* and *MYBPC3* with single affected cases as well as *TTN* and *MYOM2* with multiple affected cases (7 and 4 cases, respectively). For the four cases with *MYOM2* mutations (Fig. 1A), we showed that these genetically similar cases share similar network disturbances in gene expression (Grunert et al., 2014), which should hold true for other affected genes as well. Interestingly, the expression level of *MYOM2*-mRNA is significantly up-regulated in TOF patients with mutations compared to normal heart controls, while it appears not to be the case for patients without mutations (Fig. 1B). Two of the TOF patients with *MYOM2* mutations also harbor rare deleterious variations in *TTN* (TOF-04 and TOF-11).

**Fig. 1.**
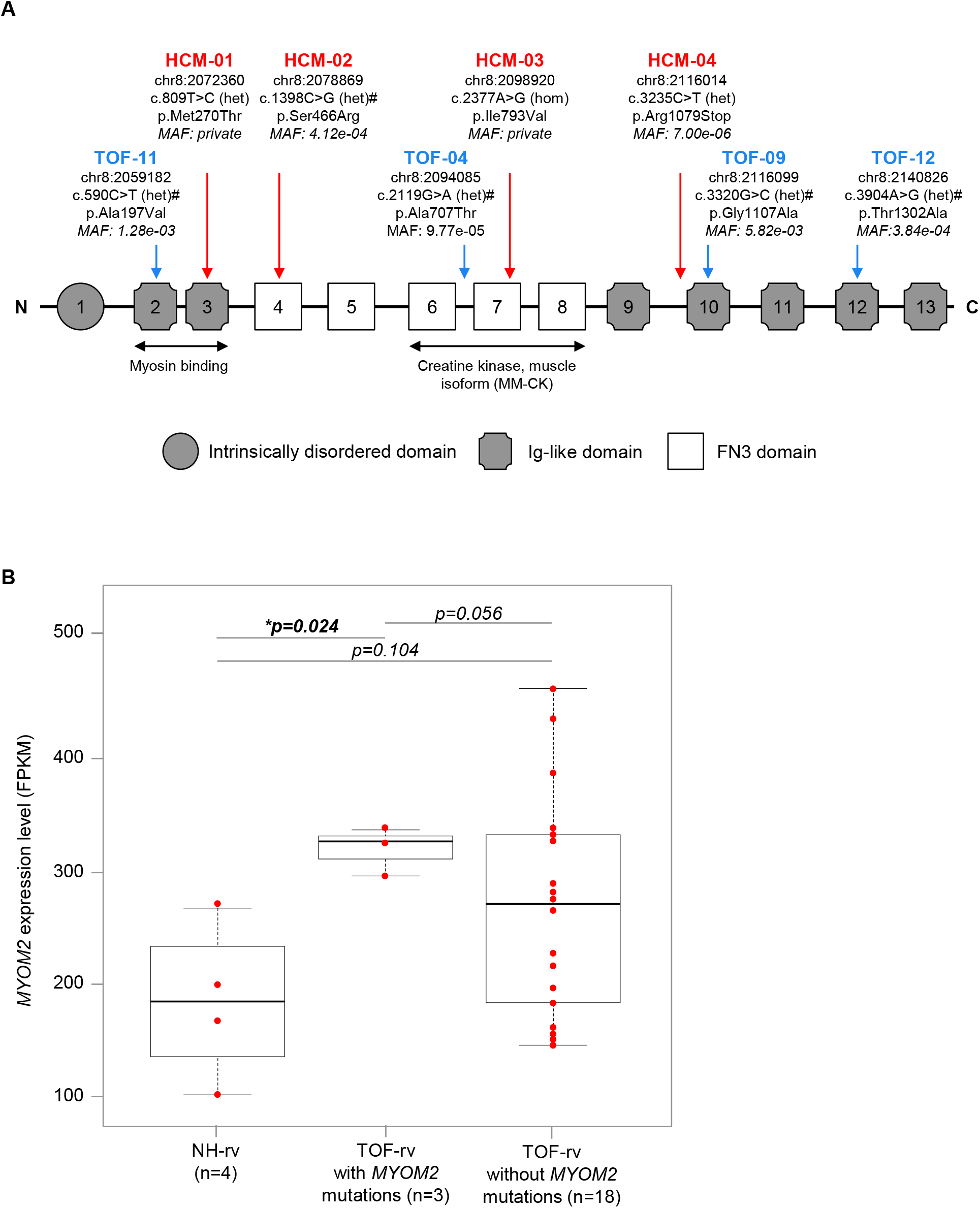
*MYOM2* mutations and expression in patients. (A) Schematic representation of MYOM2 with its domain structure and mutations found in TOF (blue) and HCM (red) patients. Mutation positions are based on human reference genome hg38. Nucleotide change based on transcript ENST00000262113. Amino acid change based on protein ENSP00000262113. Damaging prediction by PolyPhen2, SIFT or MutationTaster are indicated by ‘#’. The MAF is based on 71,702 genomes from unrelated individuals of the Genome Aggregation Database (gnomAD v3). Binding sites of two interaction partners, MYH7 and creatine kinase (muscle isoform), are indicated below. (B) RNA-seq expression level of *MYOM2* in right ventricular tissue of TOF patients and normal hearts. P-value is based on a Student’s t-test. FPKM: fragments per kilobase million; HCM: hypertrophic cardiomyopathy; het: heterozygous variation; hom: homozygous variation; MAF: minor allele frequency; NH: normal heart; RV: right ventricle; TOF: Tetralogy of Fallot. (Scheme according to Agarkova *et al*. 2005) (Agarkova and Perriard, 2005).

We further analyzed the *MYOM2* gene for putative mutations in a cohort of 66 unrelated HCM patients who had negative screening results in the known HCM disease genes (*MYH7, MYBPC3, TNNT2, TNNI3, TPM1, MYL2, MYL3, ACTC1, TCAP, TNNC1, MYOZ2* and *CSRP3*). We found four HCM patients with rare probable disease-causing mutations in *MYOM2* (Fig. 1A, Fig. S1 and Table S1). Interestingly, the mutation p.M270T is located in an Ig-like domain interacting with the light meromyosin part of the beta-myosin heavy chain, while the mutation p.I793V is located in a FN3-like domain interacting with the muscle isoform of creatine kinase (Fig. 1A). Three of the four single nucleotide variations (SNVs) are missense mutations leading to amino acid changes, while one is a truncating mutation leading to a premature stop (p.R1079X). The p.I793V mutation was the only homozygous variation. Further, we validated the initial genetic screening results in the four affected HCM patients by using a DNA resequencing array composed of known HCM disease genes. This confirmed that none of the four patients carried disease-causing mutations in the 12 most common sarcomere HCM genes. There are no *MYOM2* mutations described for HCM so far.

In general, the eight mutations found in *MYOM2* in four HCM and four TOF patients are distributed over different domains of the protein (Fig. 1A). Moreover, all variations are rare or even private as for two HCM patients (Fig. 1A), meaning they all have a minor allele frequency of less than 0.01 or zero based on 71,702 genomes from unrelated individuals of the Genome Aggregation Database (gnomAD) (Karczewski et al., 2019). Of note, almost all candidate mutations found in HCM patients of other studies occur at a very low frequency and approximately half are found in a single proband or family (Alfares et al., 2015). In addition to their very low frequency, the majority of the variations is predicted to be damaging based on PolyPhen2 (Adzhubei et al., 2013), SIFT (Vaser et al., 2016) and MutationTaster (Schwarz et al., 2010) (Fig. 1A).

### HCM-derived cardiomyocytes with MYOM2 mutation show myofibrillar disarray and reduction in passive force

Due to the severity of the LV hypertrophy, the HCM patient carrying the S466R *MYOM2* mutation (HCM-02, Fig. 1A) underwent septal myectomy (see Material and Methods for clinical courses of HCM patients). We performed morphological analysis of tissue sections and force measurements on CMs isolated from the interventricular septum of the HCM patient as well as five unaffected age-matched donors as controls.

Histological analysis showed cellular disarray and mild widening of interstitial spaces, indicative of replacement fibrosis, in the patient but not in control tissue (Fig. 2A). At the ultrastructural level, disarray of myofibrils was a frequent finding in CMs of the patient sample whereas the myofibrils of the control were regularly oriented and mostly parallel. Thus, the morphological analysis showed an HCM-dependent remodeling of the hypertrophic heart. In addition, gel electrophoretic analyses of the phosphorylation status of HCM-02 and controls showed that in particular cardiac troponin I (cTnI) was strongly phosphorylated in the controls compared to the patient (Fig. 2B and Fig. S2A). The differences might result from the usually strong protein kinase A (PKA) dependent phosphorylation in control hearts which affects especially cTnI (Kraft et al., 2013; van der Velden et al., 2003). Yet, it cannot be excluded that the *MYOM2* mutation results in additionally reduced cTnI phosphorylation in HCM-02 myocardium. Analysis of protein quantities of HCM-02 and controls suggested reduced levels of cTnI and MLC2v in HCM-02 and an increase in cMyBPC (Fig. S2B). Presumably the stoichiometry of sarcomere proteins in HCM-02 is maintained due to possible secondary effects of the MYOM2 mutation on other sarcomere proteins.

**Fig. 2.**
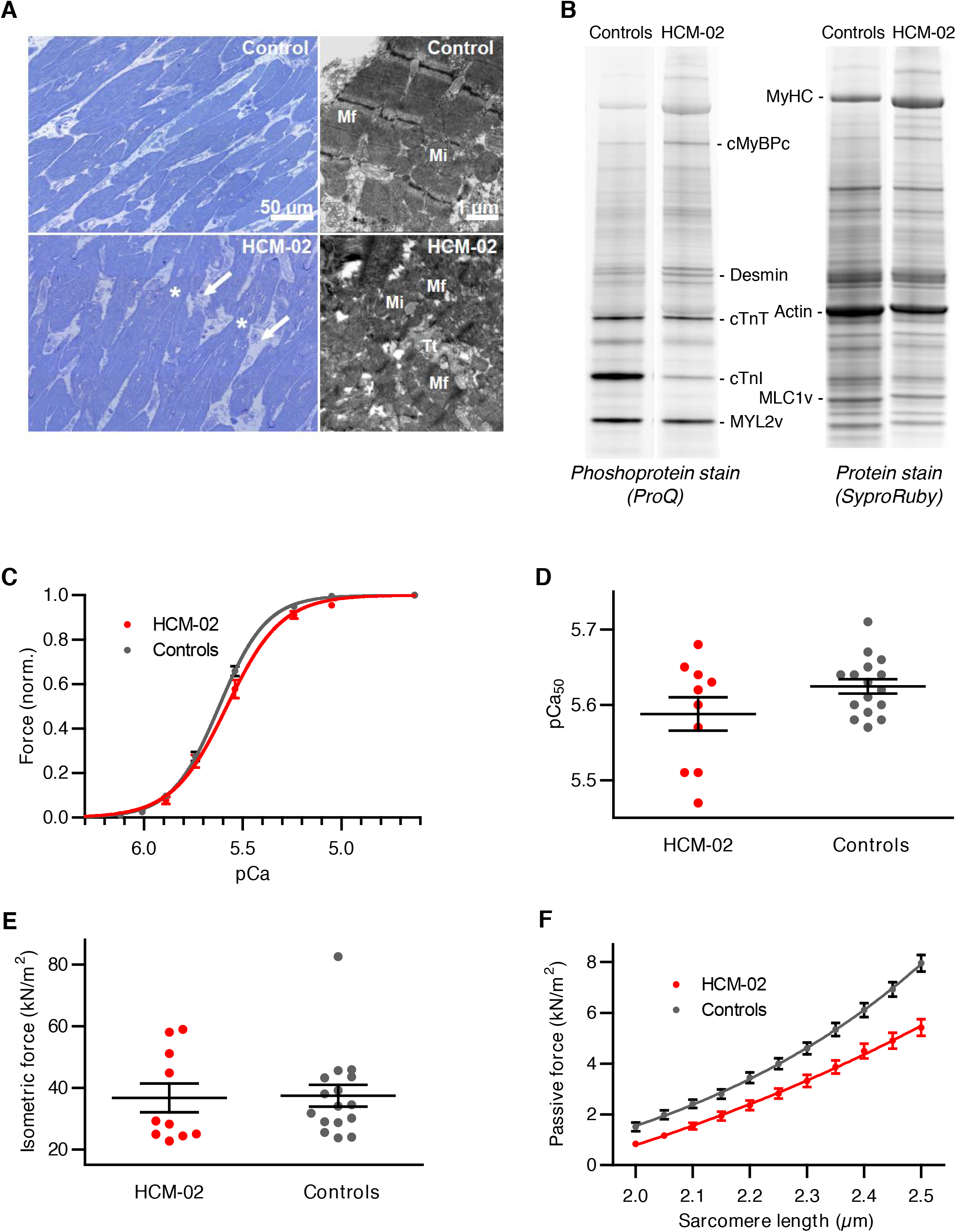
Reduced passive force in cardiomyocytes derived from HCM-patient with *MYOM2* mutation. (A) Light and electron micrographs of control (upper panel) and patient HCM-02 (lower panel) myocardial samples. In comparison to control myocardium, the patient sample showed disoriented CMs (disarray) with great variations in cellular diameter. Irregularly formed connections between CMs (asterisks) and widened interstitial spaces (white arrows) were present. Within CMs, myofibrils (Mf) of the patient frequently showed disarray with sarcomeres running in various directions whereas the sarcomeres of control tissue were mostly parallel. Mi, mitochondria; Tt, T-tubule. (B) Example for gel electrophoretic analysis of phosphorylation status of native myocardial tissue from HCM-02 and controls. For detailed analysis see Fig. S2A. (C) Isometric force generation at increasing calcium concentrations (pCa) normalized to maximum force. Lines were fitted according to a modified Hill equation (Kraft et al., 2013). (D) Calcium concentration at 50% of maximum force generation (pCa_50_) derived from fitted curves in (C). (E) Isometric force at maximum calcium activation (pCa 4.63) appeared not to be altered in HCM-02 CMs carrying the mutation. (F) Passive force at increasing sarcomere length showed a significant reduction for HCM-02 CMs carrying a *MYOM2* mutation compared to controls (p<0.01 for all sarcomere lengths). All functional analyses of CMs from HCM-02 and control in (C)-(F) occurred after adjustment of PKA-dependent phosphorylation; n=16 control CMs; n=10 HCM-02 CMs. Error bars indicate standard error of the mean.

To account for differences in phosphorylation particularly of PKA-dependent sites in cTnI (Fig. 2B), phosphorylation was adjusted by incubation of all CMs with PKA prior to force measurements. We also measured force development of CMs at different calcium concentrations and found that mutant CMs have reduced calcium sensitivity. The force-pCa curve was shifted to the right, suggesting more calcium is needed to reach 50% of maximum force development (Fig. 2C). However, the effect was not statistically significant (Fig. 2D). Force generation at maximum calcium activation was not altered (Fig. 2E).

Furthermore, we also determined passive force at increasing sarcomere lengths of CMs from patient HCM-02 and control CMs (Fig. 2F). Interestingly, passive force was significantly lower for patient CMs compared to controls at all sarcomere lengths, indicating that MYOM2 influences passive tension, in addition to TTN. Altogether, these results suggest that the mutation in MYOM2 has an effect on the passive tension of CMs which may result in altered diastolic function.

### Identification of CG14964 as putative MYOM2 Drosophila ortholog

The occurrence of *MYOM2* mutations in both, TOF and HCM patients, its altered RNA expression levels within in our TOF cohort as well as changes in the physiology of HCM-derived mutant CMs suggest an important function for the development and function of the heart. We hypothesized that it is likely to interact with critical components of the sarcomere such as MYH7, and therefore investigated any such interaction using the *Drosophila* model. *Drosophila* shares similar cardiac functions and pathways, but with lower genetic redundancy and thus, complexity compared to human or vertebrates in general (Bier and Bodmer, 2004; Bodmer, 1995; Vogler et al., 2009).

Many sarcomere proteins are conserved from human to *Drosophila*, like MYH7 (myosin heavy chain, Mhc, in flies). However, some of these only share a common domain architecture with orthology being less evident, such as *TTN* with a large number of FN3 and Ig-like domains similar to the fly genes *sallimus* (*sls*), but also to *bent* (*bt*). In the case of the myomesin family, several genes with similar domain structure comprising FN3 and Ig-like domains have been reported (DIOPT database (Hu et al., 2011)), but only a single fly gene, *CG14964*, showed a number and an arrangement of domains close to *MYOM1, MYOM2* and *MYOM3* (Fig. S3A). Of note, *CG14964* is also similar to human myosin binding proteins H (*MYBPH* and *MYBPHL)* as well as myosin binding proteins C (*MYBPC1, MYBPC2* and *MYBPC3;* Fig. S4). We suggest that in *Drosophila, CG14964* inhabits the functional space that is occupied by several human sarcomere genes, including *MYOM2* (Fig. 3A and Fig. S3B,C,D). Similarly, amino acid alignments of bt and sls to the human TTN exhibit a conserved structure of both proteins in length and regarding the tandem organization of FN3 and Ig domains (Fig. S3C,D), suggesting that both, sls and bt, adopt the function of TTN. To confirm the potential role of *CG14964* in muscle formation, we performed mRNA *in situ* hybridization (RNAscope, ACDBio) in adult fly abdomens and found *CG14964* being expressed in the heart, specifically in contractile CMs but also in somatic muscles (Fig. 3B,C and Fig. S5). Taken together, the structural conservation of the gene and cardiac expression pattern suggest that *CG14964* is the appropriate gene ortholog for studying *MYOM2* as well as other myosin binding proteins’ cardiac function in the fly.

**Fig. 3.**
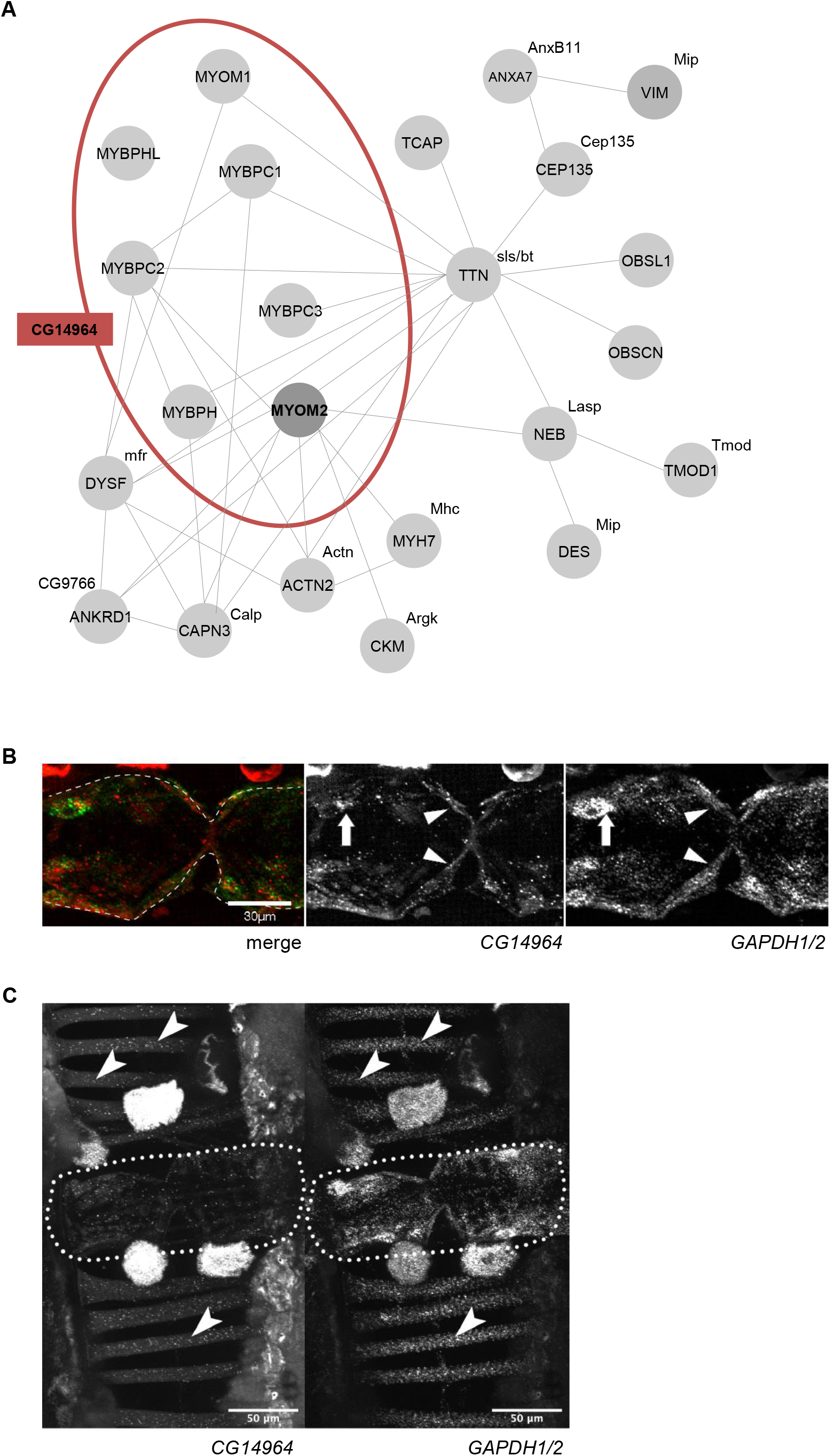
Interaction network and expression of *CG14964* - a putative ortholog of *MYOM2*. (A) Interaction network of MYOM2, TTN and MYH7 with *Drosophila* orthologs *CG14964*, bent (*bt*) and sallimus (*sls*). Physical interactions based on GeneMania v.3.6.0 (Warde-Farley et al., 2010) and other studies (Blandin et al., 2013; Brulé et al., 2010; Hornemann et al., 2003; Obermann et al., 1998). *Drosophila* orthologs based on DIOPT database (Hu et al., 2011). (B) Expression of *CG14964* in adult fly hearts (arrows/arrow heads mark perinuclear space with *CG14964* (red) and *GAPDH1/2* (green) transcript localization). Gapdh1/2 is used as a reference. (C) Expression of *CG14964* and *GAPDH1/2* in the heart (encircled), and in body wall muscles (arrows).

### Heart- and muscle-specific knockdown of CG14964 leads to heart and muscle defects

To characterize the function of *CG14964*, its expression level was reduced by RNAi-mediated knockdown (KD) specifically in the heart or in both, somatic muscles and heart, using the inducible Gal4-UAS system (Hand^4.2^-Gal4 and Mef2-Gal4, respectively) and with two independent RNAi lines were used (CG14964i-GD and CG14964i-T3 (TRiP (Perkins et al., 2015)); see Methods for further details), which both reduce *CG14964* expression in the heart and muscle measured by quantitative reverse transcription PCR (RT-qPCR) (in the heart only) and *in situ* hybridization (Fig. S6A,B,C,D). Moreover, a chromosomal deletion covering the *CG14964* locus was used alone (i.e., heterozygous deficiency that includes *CG14964*; CG14964^Df^; Fig. S6E) or with a loss-of-function allele (i.e., a homozygous CRIMIC insertion; CG14964^CRIMIC^; Fig. S6F). For both systemic mutations, CG14964^Df^ and CG14964^CRIMIC^, we characterized the KD efficiency by RT-qPCR extracted from whole flies (Fig. S6G,H,I).

To analyze the cardiac function of *CG14964*, we applied a semi-automated method to assess the contractility and rhythmicity parameters of mutant adult fly hearts using the semi-automatic optical heartbeat analysis (SOHA) method (Ocorr et al., 2014; Ocorr et al., 2009; Vogler et al., 2009). Adult flies were dissected to expose the beating heart, filmed and analyzed. In flies with moderate (∼50%) reduction of *CG14964* by RNAi or heterozygous deficient flies, we observed cardiac dilation of the beating as well as of fixed heart samples following immunostaining (Fig. 4A,B,C). Further reduction of *CG14964* by using a transheterozygous combination of a CRIMIC variant with a deficiency line resulted in cardiac constriction (Fig. 4D). A similar but not significant trend was observed with a strong KD using the TRiP line (Fig. S6B,D and Fig. S7). These data suggest that the amount of *CG14964* reduction dictates the specific heart phenotype, meaning that a moderate reduction causes dilation, whereas complete loss or strong reduction leads to restriction. Interestingly, we found that constricted hearts of *CG14964*^*CRIMIC*^ mutants show increased expression of Mhc (Fig. S9), indicating that reduction of *CG14964* might cause myofibrillar hypertrophy in a dosage-dependent manner.

**Fig. 4.**
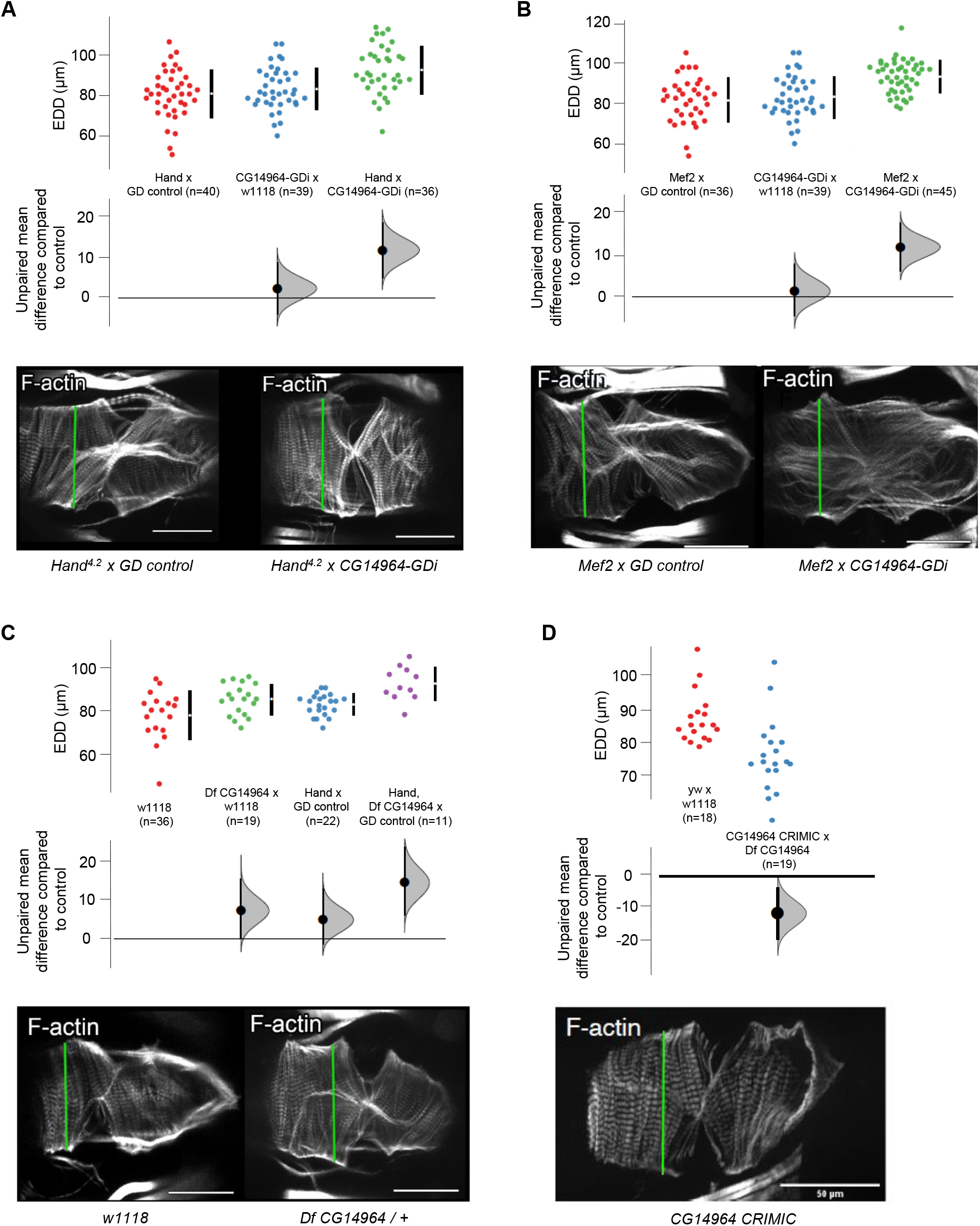
Cardiac-specific knockdown of *CD14964* leads to dosage-dependent heart defects in the adult fly. End-diastolic diameter (EDD) of 3 weeks old flies harboring a knockdown of *CG14964* by using (A) *CG14964*-GDi crossed with *Hand*^*4*.*2*^-Gal4, (B) CG14964-GDi crossed with *Mef2*-Gal4, (C) *CG14964* deficiency or (D) transheterozygous *CG14964*^*CRIMIC*^/deficiency. We observed heart dilation (A-C, mild knockdown) or constriction (D, strong knockdown). All raw data are shown (with mean and standard deviation) as well as effect size and 95%-CI below the data. In addition, Phalloidin-stained cardiac myofibrils show altered heart diameters in representative examples (measurements taken at green line). Scale bar = 50μm.

As *CG14964* is expressed in somatic muscle, we further examined the role of *CG14964* for muscle function in general. Using the Rapid Iterative Negative Geotaxis (RING) assay (Gargano et al., 2005) on flies with Mef2-Gal4 mediated KD, we found a significant locomotion (i.e., climbing) defect in 3 weeks old CG14964i-T3 flies compared to CG14964i-GD and control flies (Fig. 5A). Similarly, muscle- and heart-specific KD of *CG14964* using CG14964i-T3 also resulted in a reduced lifespan of adult flies: the half-survival rate of CG14964i-T3 flies was severely decreased compared to CG14964i-GD and control flies, with less than half of the flies surviving beyond week 4 (Fig. 5B). The reduced climbing ability together with a decreased lifespan of muscle-specific KD of *CG14964* of adult flies indicate an impairment of the overall muscle function, which implies that *CG14964* seems to be required in all somatic muscles. This is in line with our observation that *CG14964*^*CRIMIC*^ mutants either homozygous or *in trans* to the *CG14964* deficiency also showed severely compromised climbing abilities and reduced viability in addition to cardiac defects. Lastly, we also tried to overexpress human MYOM2 in *Drosophila* but while we could detect the human transcript when driving the hMYOM2 cDNA with Mef2-Gal4 (Fig. S10) we were not successful in obtaining evidence for hMYOM2 by antibody staining or mScarlet localization, indicating that protein translation and or maturation of hMYOM2 in fly tissue is not trivial.

**Fig. 5.**
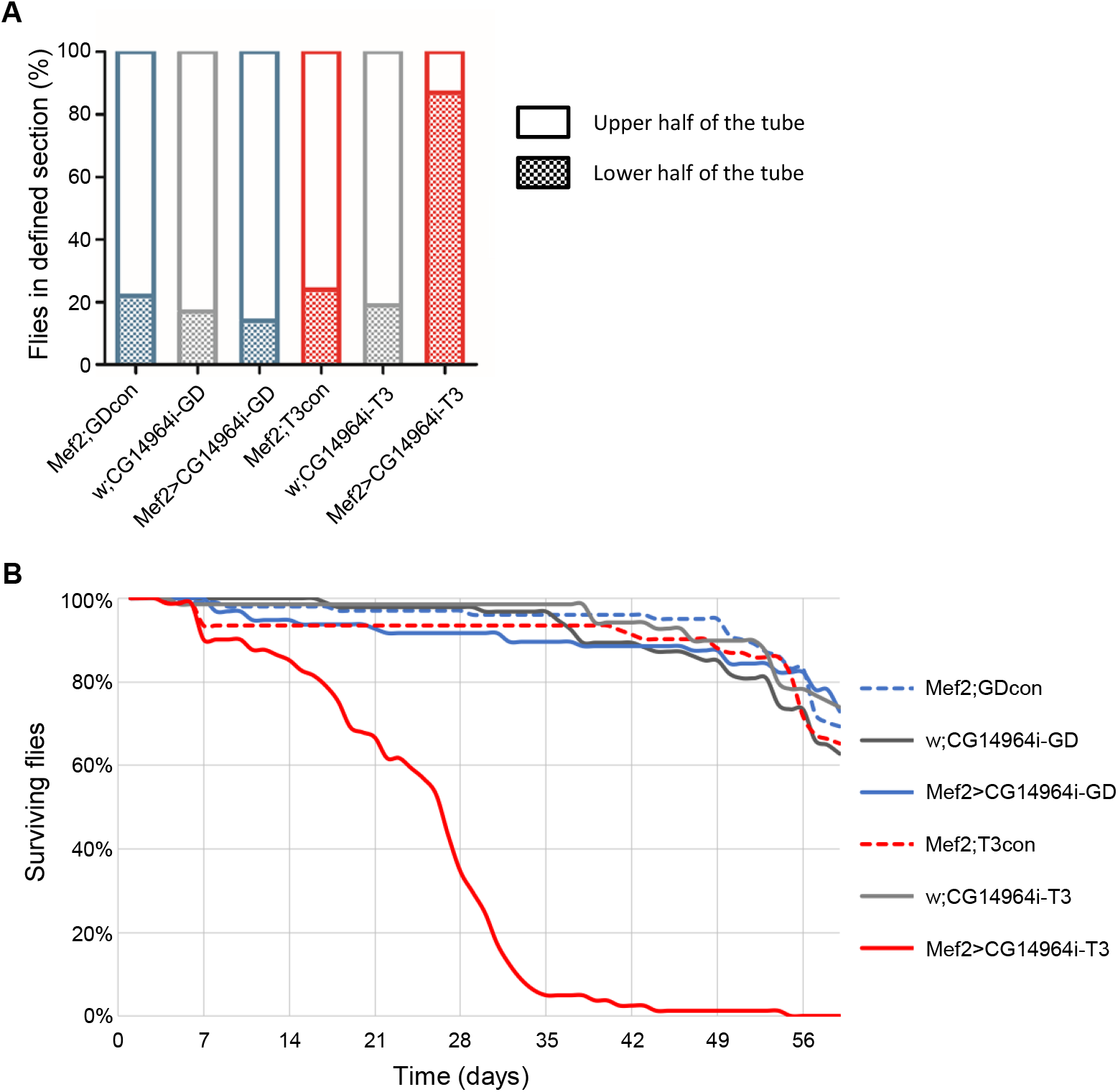
Muscle-specific knockdown of *CG14964* causes locomotion defects and reduced lifespan in adult flies. (A) Locomotion test performed using the RING assay on adult flies expressing *CG14964i*-GD (left) and *CG14964i*-T3 (right) in muscles (*Mef2*-Gal4) showed reduced locomotion ability at 3 weeks in *Mef2*>*CG14964i*-T3 flies only. Graph shows percentage of fly population in defined section of the vial after 20 sec. (B) A survival assay performed on *Mef2*>*CG14964i*-GD and *Mef2*>*CG14964i*-T3 with appropriate controls revealed a decreased survival for *Mef2*>*CG14964i*-T3 with less than half flies surviving 28 days versus more than 90% in the other tested lines.

### Interaction to Mhc (MYH6/MYH7)

Since it was shown that MYOM2 physically interacts with MYH7 *in vitro* (Obermann et al., 1998), we wanted to test whether *CG14964* and myosin-heavy chain (*Mhc*), the fly ortholog for *MYH6* and *MYH7*, could interact at the genetic level *in vivo*. We used a heterozygous null mutant for *Mhc* (*Mhc*^*1*^) (O’Donnell and Bernstein, 1988; Wells et al., 1996) combined with different KD lines for *CG14964* (both GD and TRiP). Heterozygosity for *Mhc*^*1*^ alone causes a modest but significant cardiac constriction (Fig. 6; far right orange data points). Reduction in *CG14964* levels using the strong TRiP line (Fig. 6A; middle green data points) or using the moderate GD line (Fig. 6B; middle green data points), causes a decrease or an increase in heart diameter, respectively, compared to controls (Fig. 6; left data points). Strikingly, the combination of *Mhc*^*1/+*^ with muscle-specific *CG14964* KD results in a different interaction depending on the RNAi alone, meaning *Mhc*^*1/+*^ combined with CG14964i-T3 results in a further constriction, compared to *Mhc*^*1/+*^ alone (Fig. 6A; purple data points), indicating a synergistic enhancement of constriction. Similarly, heterozygosity for *CG14964*^*CRIMIC*^ also reduces heart diameter in an *Mhc*^*1/+*^ background (Fig. S8). In contrast, *Mhc*^*1/+*^ combined with *CG14964i*-GD results in intermediate cardiac diameters, compared to either one alone (Fig. 6B; purple data points), indicating a normalization to wildtype. This means that the dilation phenotype of CG14964i*-GD* is reversed by the reduction of *Mhc*, which is consistent with our observation of significantly increased Mhc levels in a *CG14964* mutant background. Taken together, our data suggest that *CG14964* and *Mhc* functionally interact, consistent with the postulated equivalent functions of *MYOM2*-*MYH6/7* in mammals and *CG14964-Mhc* in the fly heart.

**Fig. 6.**
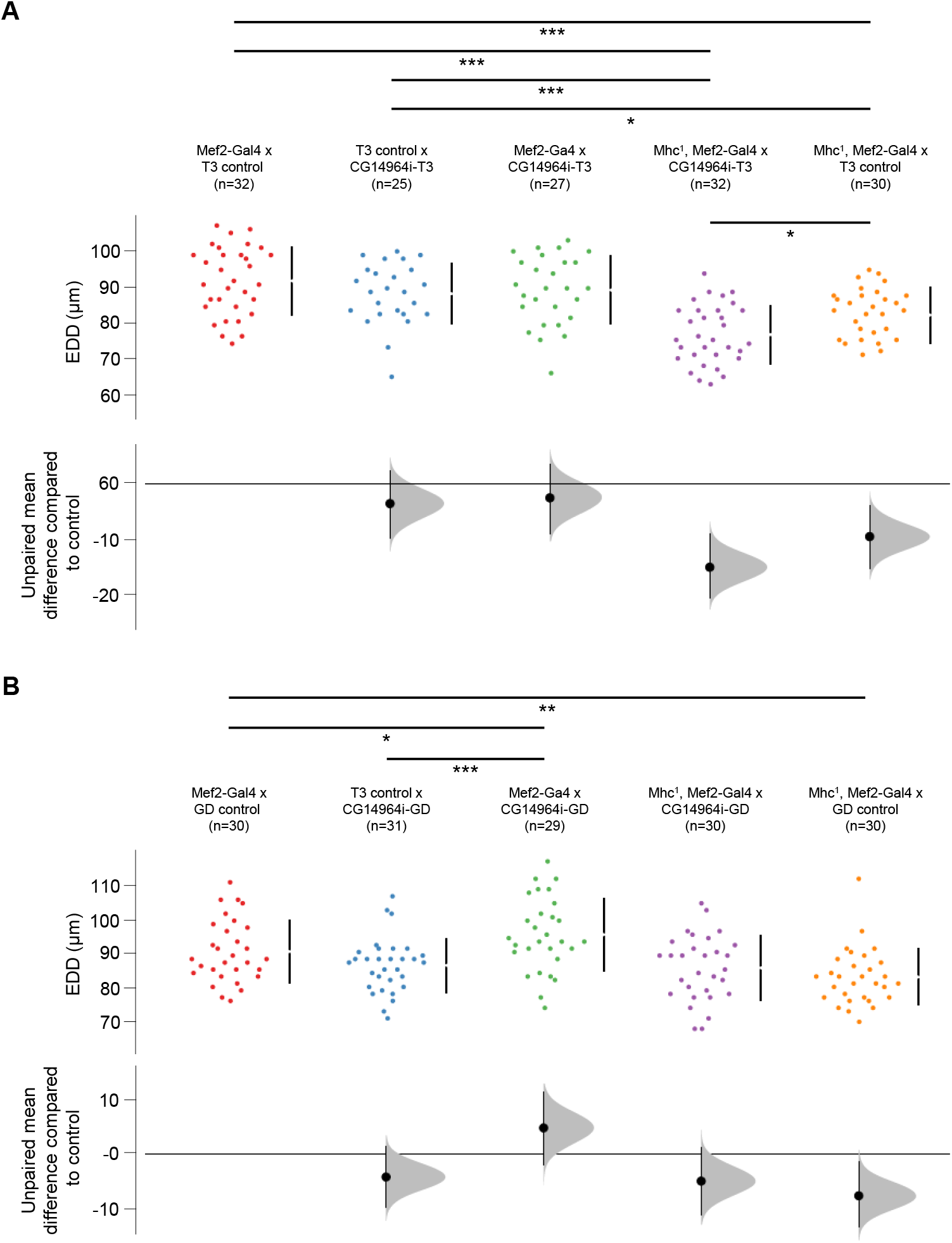
Interaction between *CG14964* and *Mhc*. (A) End-diastolic diameters (EDD) are decreased in *Mhc*^*1*^ heterozygous flies (orange) and become further constricted upon strong *CG14964* knockdown (purple). (B) Mild *CG14964* knockdown causes enlarged hearts (green), which is restricted by *Mhc*^*1*^ heterozygosity (purple). All raw data are shown (with mean and standard deviation) as well as effect size and 95%-CI below the data. For *Mhc - CG14964* interaction, significance was tested using Student’s t-test (*p<0.05; **p<0.01; ***p<0.001).

## DISCUSSION

Myofibrils mediate skeletal and cardiac muscle contraction in vertebrates and invertebrates. Their basic unit is the sarcomere with two transverse structures, the Z-disk and the M-band, which anchor thin (actin) and thick (myosin) filaments to an elastic system (Lange et al., 2020). Alterations of sarcomere proteins, therefore, influence the contractile performance of the heart and skeletal muscle (Hamdani et al., 2007). In this study, we found rare deleterious genetic variations in the *MYOM2* gene of HCM and TOF patients, a well-characterized protein of the sarcomere and a major structural component of its myofibrillar M-band. In humans, there are besides *MYOM2*, which is specifically expressed in the heart and skeletal muscle, two other myomesin genes, namely *MYOM1* and *MYOM3*. However, this apparent redundancy in myomesins does not result in a buffering effect *per se*, which can be seen for example in the genomics of Arthrogryposis (Pehlivan et al., 2019). Here, loss-of-function of *MYOM2* can result in termination of gestation of the affected fetus with cardiac and arthrogryposis findings, without variation in *MYOM1* or *MYOM3* (Pehlivan et al., 2019).

We observed four affected cases with *MYOM2* mutations in our small, but phenotypically very homogeneous cohort of 13 isolated TOF patients (∼31%). Moreover, these cases share differential expression profiles in other mutated genes, including *MYOM2*, and exhibit an aberrant histological phenotype of the right ventricular tissue (Grunert et al., 2014). In a recent study comprising more than 2,800 CHD patients (PCGC study), only 14 cases with rare inherited or *de novo* mutations in *MYOM2* were found, of which two (14%) are TOF patients (Jin et al., 2017). As different as in many CHD cohorts in general, the variety of CHDs with *MYOM2* mutations in the PCGC study is very wide, comprising complex heterotaxias such as double outlet right ventricle as well as isolated simple atrial and ventricular septal defects (Jin et al., 2017). However, there are subgroups of TOF patients, which in the case of the PCGC study comprise a relatively small number of cases and for our small but homogeneous group of patients, in terms of their clinical parameters and features, a relatively high number of cases with *MYOM2* mutations. These mutations might contribute to the phenotype during heart development (Grunert et al., 2014) as well as in the long-term. The latter is reflected by an arrhythmic burden in adult TOF patients (Khairy et al., 2010); (Wu et al., 2014) and that the majority of mutated genes show continuous expression during adulthood (Grunert et al., 2014).

It has been shown that TOF is genetically heterogeneous and a subgroup is characterized by genetic alterations in other sarcomere genes known to be causative of cardiomyopathy such as *MYH7* and *MYBPC3* (Grunert et al., 2014; Vermeer et al., 2016). Thus, we screened for *MYOM2* mutations in a cohort of 66 HCM patients that had no mutations in the known disease genes. Our screening revealed a subgroup of four HCM patients with rare *MYOM2* mutations, half of which predicted to be damaging. In general, the genetic testing in HCM was limited by contemporary technologies. However, we have validated the screening approach using an Affymetrix resequencing array as an up-to-date and sensitive method. Approximately 40% of HCM cases have a non-familial subtype (Ingles et al., 2017), however since data to estimate co-segregation within families was not available we cannot exclude this possibility. Furthermore, it may be possible that referral bias may have led to an overestimate of the frequency of *MYOM2* mutation carriers in HCM because of the retrospective nature of this study. However, very few data exist concerning a pathogenic role for mutations in M-band proteins, especially the myomesin protein family, in cardiomyopathies so far. The first study suggesting a link described the missense mutation p.V1490I in *MYOM1*, which co-segregated with the phenotype in a family with HCM (Siegert et al., 2011). In a later study, a panel of 62 sarcomere and non-sarcomere genes (including *MYOM1* but not *MYOM2*) in 41 HCM patients was investigated by high-throughput sequencing. In total, they found three rare *MYOM1* SNVs, two with and one without deleterious prediction (Bottillo et al., 2015). Furthermore, a panel of 108 cardiomyopathy and arrhythmia-associated genes including only *MYOM1* was screened in 24 patients with restrictive cardiomyopathy (Kostareva et al., 2016). Here, two variants of unknown significance (VUS) were identified in *MYOM1*. Two further *MYOM1* VUS were found in one young sudden death victim with cardiac dilatation by whole exome sequencing-based molecular autopsy (Shanks et al., 2017). Two further studies screened patients with dilated cardiomyopathy (Akinrinade et al., 2015; Marston et al., 2015). Akinrinade *et al*. used a targeted resequencing panel of 101 genes (including *MYOM1* and excluding *MYOM2*) for screening of 145 Finnish patients and identified one SNV in *MYOM1* (designated as VUS) in two individuals (Akinrinade et al., 2015). Of note, the study by Marston *et al*. investigated a cohort of 30 patients with familial dilated cardiomyopathy in 58 cardiomyopathy-related genes. Indeed, it is the only study including *MYOM2* besides *MYOM1* (Marston et al., 2015). However, they found a SNV in *MYOM1* while no rare SNVs in *MYOM2* were detected. Other larger studies screening different defined gene panels (from 31 to up to 126 genes related to heart disease) by targeted resequencing in cardiomyopathy patients did not include *MYOM1* and *MYOM2* into their panels (Cuenca et al., 2016; Haas et al., 2014; Lopes et al., 2013; Walsh et al., 2017). In summary, while one probably disease-associated mutation as well as seven VUS have been identified in *MYOM1*, no or no rare *MYOM2* variations in cardiomyopathy patients have been described so far.

Functional insight into the role of *MYOM2* came from morphological analysis of tissue sections and force measurements on CMs isolated from the interventricular septum of a patient that underwent myectomy (HCM-02). Interestingly, the HCM-derived CMs showed significantly reduced passive force, indicating that the *MYOM2* mutation influences the passive properties of the patient-derived CMs. This is an interesting effect, which could indicate that, in addition to titin, Myomesin-2 also has an influence on the diastolic properties of the CMs and the ventricle. This observation would be in line with the role of myomesin as part of the shock absorber function of the striated muscle M-band that together with titin stabilizes the myosin filament lattice longitudinally (Lange et al., 2020; Schoenauer et al., 2005).

In addition, we studied the potential cardiac and muscle function of MYOM2 and the interaction partner MYH7 by assessing the effect of down-regulation in *Drosophila. Drosophila* is an excellent animal model for studying the genetics of human disease mechanisms, which is typically masked due to genetic redundancy present in mammals (Pickett and Meeks-Wagner, 1995). There is no current mammalian animal model for *MYOM2*, and in zebrafish, four redundant paralogous myomesin genes exist of which only MYOM3 was studied in detail (Xu et al., 2012). Given the advantages of the fly model, we probed the function and genetics of *MYOM2* by analyzing the closest functional ortholog *CG14964* (*dMnM)* in *Drosophila*, a gene that is similar to myomesins as well as other myosin binding proteins. Cardiac-specific knockdown or deletion of *CG14964* leads to dosage-dependent heart defects (i.e., cardiac dilation or constriction) and the muscle-specific knockdown causes locomotion defects and reduced lifespan of adult flies. The larger end-diastolic diameter in *Drosophila* suggests either mild (eccentric) hypertrophy or reduced stiffness of the myofibrils/sarcomeres upon *CG14964* KD. This is consistent with reduced passive force at different sarcomere length and thus, reduced stiffness observed in human CMs with mutation Ser466Arg (HCM-02) in *MYOM2*. In addition, it supports the hypothesis that *CG14964* is a myomesin ortholog in *Drosophila*. The differences in the knockdown results may be explained by their efficiencies, given that the strongest KD phenotype was comparable to the combination of two strong alleles *in trans*. Of note, the cardiac phenotype of *CG14964 RNAi* appears to be dose-dependent, with mild reduction causing cardiac dilation that is reversible by reduction in Mhc dosage, while strong reduction leads to cardiac constriction potentially as a result of excessive increase in myosin. This is reminiscent of specific mutations in MYH7 (encoding ß-myosin heavy chain) in humans that can give rise to, for example, dilation or restriction (Hershberger et al., 2013). In addition to classic HCM, MYH7 mutations may cause cardiomyopathies with different heart morphology and function such as dilated cardiomyopathy (DCM) and HCM with features of restrictive *cardiomyopathy* (RCM) (Bondue et al., 2018; Kubo et al., 2007; Møller et al., 2009). Indeed, the cardiac phenotype caused by MYH7 mutations shows a great variety ranging from late onset DCM with mild to moderate dilatation (Villard et al., 2005) to severe pediatric RCM (Ware et al., 2007). Similarly, in *Drosophila*, different types of *Mhc* mutations with inhibited or increased motor activity show *cardiac* dilation or constriction, respectively (Cammarato et al., 2007; Kronert et al., 2018). However, in contrast to MnM loss of function where we do not see sarcomere defects, strong knockdown of Mhc causes breakdown of the sarcomere. Therefore, we hypothesize that the level of MnM is important to finetune sarcomere function potentially by regulating Mhc levels, but not for the overall structure of the sarcomere.

*MnM (CG14964)* seems to be a hub gene within the interaction network of sarcomere genes as shown in Fig. 3A. Four interaction partners of *MYOM2* are known to carry mutations causing cardiomyopathies (*MYH7, ANKRD1, ACTN2, TTN*), while two are known to cause muscular dystrophy (*DYSF, CAPN3*). As 30-50% of HCM cases harbor mutations in the sarcomere gene *MYH7* (Vermeer et al., 2016), we tested and showed synergistic interaction of *CG14964* with *Mhc* (*MYH6*/*MYH7*) in the adult fly.

In summary, this study showed novel rare, probably disease-relevant mutations in the sarcomere gene *MYOM2* of patients with TOF and HCM, in particular for cardiomyopathy patients. Moreover, the functional characterization of affected patient-derived CMs as well as functional analyses of the up-to-now unknown fly gene *CG14964*, as the likely ortholog of *MYOM2* as well as other myosin binding proteins, suggest that *MYOM2* is involved in development of the heart and plays critical roles in establishing or maintaining robust heart function. Thus, *MYOM2* is a disease candidate gene for HCM and TOF, both of which exhibit an impaired left and right ventricular function, respectively.

## MATERIALS AND METHODS

### Phenotyping of HCM patients

The local institutional review board of the Charité – Universitätsmedizin Berlin approved the study and written informed consent was obtained from all participants. The study protocol conforms to the ethical guidelines of the 1975 Declaration of Helsinki.

The study cohort comprised 66 unrelated patients with HCM of German origin (29 % female and 71% male). All patients had negative screening results in known HCM disease genes (see below). While 20 patients showed a non-obstructive form of the disease (about 30%), 46 patients (70%) had hypertrophic obstructive cardiomyopathy (HOCM) presenting with left ventricular outflow tract obstruction at rest and/or with exercise (often accompanied with systolic anterior motion of the anterior mitral leaflet). This distribution is in accordance with the general distribution in HCM (Maron et al., 2006). The cohort was examined on the basis of medical history, physical examination, 12-lead electrocardiogram (ECG), and two-dimensional and M-mode echocardiography. As clinically indicated, cardiac magnetic resonance imaging, heart catheterization, angiography, and Holter ECG were performed in some patients. The diagnosis of HCM was made according to the established criteria (Elliott et al., 2007; Gersh et al., 2011). The pathological hallmark of HCM is unexplained left ventricular hypertrophy. Briefly, the major inclusion criterion was the presence of an interventricular septal thickness (IVS) ≥ 13 mm (15 mm if sporadic, 13 mm if familial) in the absence of loading conditions (hypertension, valve disease) sufficient to cause the observed abnormality and in the absence of systemic disease such as amyloidosis.

### Genetic analysis in HCM patients

The frequent HCM disease genes *MYH7* and *MYBPC3* as well as rare disease genes (*TNNT2, TNNI3, TPM1, MYL2, MYL3, ACTC1, TCAP, TNNC1, MYOZ2, CSRP3*) were screened as previously published by us and colleagues (Geier et al., 2003; Kabaeva et al., 2002; Mogensen et al., 2004; Perrot et al., 2006; Perrot et al., 2005; Posch et al., 2008). Disease-causing mutations in all these genes were excluded in the 66 HCM patients comprising the study cohort (see above). For validation purposes, randomly selected 13 patients (20% of the cohort) were additionally analyzed by a custom DNA resequencing array composed of eleven HCM disease genes (Affymetrix) as previously described (Fokstuen et al., 2008; Fokstuen et al., 2011). In addition, the four HCM patients who carried a *MYOM2* mutation (Table S1) were also analyzed using this technique. Rare probable disease-causing mutations in HCM genes were identified neither in the 13 patients nor in the *MYOM2* mutation carriers. Further, mutations in *MYOM1* were also excluded in these four patients.

*MYOM2* screening was performed using Sanger sequencing. Briefly, the 36 coding exons of *MYOM2* were PCR-amplified using flanking intronic primers and directly sequenced using ABI Big Dye Terminator chemistry. Found variants were annotated according to the cDNA and protein reference sequence (Genbank ID NM_003970.3, Ensembl ID ENST00000262113.8, and UniProtKB/Swiss-Prot P54296). The significance of the found variants was further analyzed in a first step by considering the nature and location of the change and its frequency found in large population-based datasets (such as from the Exome Aggregation Consortium (Lek et al., 2016) and the Genome Aggregation Database (Karczewski et al., 2019)) and own controls (minor allele frequency below 0.02% as recommended by Burke *et al*. for cardiomyopathies (Burke et al., 2016)). In a second step, the conservation of the affected amino acid and the possible functional impact of the variants (using mutation prediction tools such as PolyPhen2 (Adzhubei et al., 2013), SIFT (Vaser et al., 2016) and MutationTaster (Schwarz et al., 2010) were assessed.

### Clinical courses of HCM patients with MYOM2 mutations and control heart tissue

All four unrelated, non-familiar cases were characterized by left ventricular (LV) hypertrophy and electrocardiogram abnormalities (see detailed clinical data in Table S1). They showed a symptomatic form of hypertrophic obstructive cardiomyopathy (HOCM) which led to an invasive septal reduction intervention in two of them. All patients were of European (German) ancestry.

The female patient HCM-01 (carrier of mutation p.Met270Thr) presented with symptoms such as angina pectoris, dyspnea on exertion, dizziness, and palpitations at the age of 50 years (New York Heart Association (NYHA) functional class II). Onset of disease was at the age of 31 years. She was found to have hypertrophic cardiomyopathy with outflow tract (OFT) obstruction(interventricular pressure gradient at rest of 20 mm Hg measured during cardiac catheterization). Echocardiography revealed an interventricular septal (IVS) thickness of 16 mm and a posterior wall (PW) thickness of 12 mm; ejection fraction (EF) was 35%. The patient was treated with calcium antagonists which improved symptoms.

The male patient HCM-02 (carrier of mutation p.Ser466Arg) had an early onset of disease at the age of 19 years. Later on, he developed a symptomatic form with angina, dyspnea on exertion, and syncopes (NYHA class III). He showed a moderate LV hypertrophy with an IVS of 15 mm and with a PW of 14 mm as well as normal cardiac dimensions at the age of 52 years. Heart catheterization confirmed the diagnosis of HOCM with an OFT gradient of 100 mm Hg at rest. Because of this OFT obstruction (including systolic anterior motion (SAM) of the mitral valve and mitral insufficiency), the53-years-old patient underwent surgical septal myectomy (Morrow procedure) which released the OFT obstruction. Three years later, an automated implantable cardioverter/defibrillator (AICD) was implanted because of ventricular tachycardia accompanied with recurrent syncopes. Because of atrial flutter, a cavotricuspid isthmus ablation was performed at the age of 68 years.

The male patient HCM-03 (carrier of mutation p.Ile793Val) had disease onset at the age of 50 years. He complained about dyspnea and dizziness (NYHA class II) and the diagnosis of HOCM was made at the age of 56 years. He had a resting OFT gradient of 40 mm Hg (measured during catheterization) including SAM. Echocardiography revealed an LV hypertrophy with an IVS of 17 mm and a PW of 14 mm as well as normal cardiac dimensions; his fractional shortening (FS) was also normal (40-42%). Under high dose calcium antagonist treatment, his OFT gradient was below 10 mm Hg and his symptoms improved.

The female patient HCM-04 (carrier of mutation p.Arg1079X) had a late onset of disease at 60 years of age. Hypertrophic cardiomyopathy was diagnosed at the age of 69 years. At this age, she showed symptoms such as palpitations, syncope, angina, and dyspnea (NYHA class III). Echocardiography showed severe LV hypertrophy with an IVS of 23 mm and a PWT of 16 mm as well as normal FS of 42%. Because of an OFT gradient of 95 mm Hg at rest, the patient underwent two subsequent transluminal septal ablations (an embolization of the first and second septal branches) at the age of 72 years. These led to a reduction of the gradient and improvement of symptoms but LV hypertrophy was just slightly reduced (as shown by subsequent echocardiography measurements). About two years after septal ablation, symptoms worsened again and also the pressure gradient raised to 50 mm Hg. Further, the patient developed intermittent atrial fibrillation.

As control tissue, we used flash frozen tissue from the interventricular septum of five unaffected, age-matched non-transplanted donor hearts (3 males and 2 females of age 23-56 years) from Sydney Heart Bank (for SHB heart codes and details see Table S2).

### Morphological analyses

Flash frozen samples (HCM-02 and control) were slowly thawed on melting fixative (1.5% paraformaldehyde, 1.5 % glutaraldehyde in 0.15 M Hepes buffer). After aldehyde fixation, samples were subsequently postfixed with osmium tetroxide, bloc-stained with uranyl acetate, dehydrated in acetone and embedded in epoxy resin. Semi-und ultrathin sections were cut and stained with toluidine blue and lead/uranyl, respectively.

### Force measurements, and phosphorylation and protein analyses of CMs from HCM-02 and control myocardium

Protein analysis and force measurements on isolated CMs from heart tissue (interventricular septum) were performed for one HCM patient (HCM-02; male, 53 years) as well as five donor hearts serving as controls (on average 38 years). Written informed consent for use of the tissue and approval of the local ethics committee of Hannover Medical School for the study on human tissue was obtained (No. 507/09). For analysis of phosphorylation of cMyBPC, cTnT, cTnI, and MLC2v in native myocardial tissue from HCM-02 and controls (Fig. S2A) gradient gels were used and phosphorylation was analyzed by calculating the ratio of Pro-Q Diamond staining (phosphorylated protein) vs. SYPRO Ruby staining (total amount of protein) for each band, respectively, as described elsewhere (Kraft et al., 2013). To study relative protein quantities of cMyBPC, cTnT, cTnI, and MLC2v in HCM-02 and control tissue (Fig. S2B), the bands of the respective proteins on the SYPRO Ruby stained gels (Fig. 2B) were analyzed densitometrically and normalized to the α-actinin signal in the same lane.

For functional studies, CMs were isolated and force was measured as described in detail elsewhere (Montag et al., 2018). Briefly, CMs were mechanically isolated from myocardial tissue and chemically permeabilized with triton-x-100. Single CMs were mounted between force transducer and lever arm of a custom-made setup for biomechanical characterization. Active force was measured by incubating CMs in physiological solutions with different calcium-concentrations. Passive force was determined in relaxing solution (pCa 8.0) by applying step release and re-stretch of the CMs (Montag et al., 2018) at increasing sarcomere lengths starting from 1.95 µm up to 2.4 µm. All measurements occurred after incubating the CMs with protein kinase A (PKA) to adjust PKA-dependent phosphorylation of sarcomeric proteins (Kraft et al., 2013). 16 and 10 CMs were measured for controls and HCM-02, respectively.

### Fly husbandry

All fly stocks were maintained at 25°C on standard fly food medium. The following fly stocks were used: w^1118^ (#3605, Bloomington Drosophila Stock Center (BDSC)), GD control (#60000, Vienna Drosophila Resource Center (VDRC)) and TRIP control (#36303, BDSC) as control flies, DMef2-Gal4 (Ranganayakulu et al., 1996) and Hand^4.2^-Gal4 (Han and Olson, 2005) as driver lines. For *CG14964*: Df(3L)BSC672 (#26524, BDSC), CRIMIC line CG14964^CR01157-TG4.1^ (#81199, BDSC); *CG14964* RNAi lines: GD (#43603, VDRC) and TRIP3 (#65245, BDSC). For *Mhc*, we used *Mhc*^*1*^ mutant (O’Donnell and Bernstein, 1988) and *Mhc*^*YD0783*^ (#50881, BDSC).

### Heart functional analysis

To assess heart function in adult flies, we used the Semi-automated Optical Heartbeat Analysis (SOHA) method (Cammarato et al., 2015; Fink et al., 2009; Ocorr et al., 2009). Briefly, adult flies were anesthetized using FlyNap (#173025, Carolina) and dissected in artificial hemolymph to expose the dorsal heart (Vogler and Ocorr, 2009). Prior to filming, the hearts were allowed to equilibrate with oxygenation for 15–20 minutes. 30-second movies were recorded with a Hamamatsu Orca Flash 4.0 camera (at 140 frames/sec) using a Zeiss A1 Axioscope (10x magnification). Movie analysis was performed by SOHA software (Oaktree Technologies, www.sohasoftware.com) (Cammarato et al., 2015).

### Overexpression of hMYOM2 in Drosophila

A full-length cDNA of hMYOM2 was obtained (MGC Human MYOM2 Sequence-Verified cDNA (CloneId:6205359, Dharmacon) and cloned into pUASattB (Bischof, 2007). For N- or C-terminal tagging of hMYOM2 using Gibson assembly, mScarlet was amplified from a plasmid template using overlapping primers to facilitate in-frame assembly into hMYOM2-pUASattB. Constructs were inserted into attP2 using a commercial injection service (Bestgene, Inc.).

### Climbing assay

Climbing defects were quantified using the Rapid iterative negative geotaxis (RING) assay which was performed as previously described (Gargano et al., 2005), with the following changes: adult flies were transferred to an empty fly tube and let to adapt for 10 min. The tubes were tapped 3 times to trigger the negative geotaxis response and 30 sec intervals were recorded to document the distance the flies could climb up. This experiment was performed in triplicates for each biological replicate and the mean of these 3 replicates was calculated for each experiment.

### Survival assay

The measurement of adult lifespan in Drosophila was performed as previously described (Linford et al., 2013). Briefly, control and experimental female flies were collected by CO2 anesthesia and transferred to tubes with fresh standard food on Day 0 (D0). 10 vials, (containing 10-15 flies) were collected per genotype (100-150 flies in total). Flies were transferred to a new tube with fresh food every 2 days and the number of dead flies was counted. This was repeated until the last fly died.

### RNA isolation and RT-qPCR

Total RNA was isolated from about 15-20 adult fly hearts, using TRIzol reagent (Invitrogen) combined with the Quick-RNA MicroPrep Kit (Zymo Research), including a step of DNAse-on-column treatment, following manufacturer’s instructions. RNA quality and quantity were respectively assessed using Agilent RNA 6000 Pico kit on Agilent 2100 Bioanalyzer (Agilent Technologies) and Qubit RNA HS assay kit on Qubit 3.0 Fluorometer (Thermo Fischer Scientific). Total RNA was reverse-transcribed using the PrimeScript RT Master Mix (Takara). RNA from 3-5 adult female whole flies was isolated using TRIzol reagent combined with Chloroform/Ethanol extraction. RNA quality and quantity were respectively assessed using Nanodrop spectrometer. cDNA was generated using Superscript IV Reverse Transcriptase (Invitrogen) with additional DNase I treatment or using QuantiTect Reverse Transcription Kit (QIAGEN). SYBR Green based real-time qPCR (Sybr Green I Master, Roche) was performed on a LightCycler480 (Roche) and LightCycler96 (Roche). Gene expression quantification was determined using the 2^-ΔΔCT^ method (Pfaffl, 2001), with *Rp49* as a reference gene. Values are derived from 3-5 biological replicates.

### Immunostainings and fly heart measurements

The immunostaining of fly adult hearts was performed as described previously (Alayari et al., 2009). Fly hearts were dissected as described for the SOHA method (see above), and myofibrils were relaxed using 10mM EGTA followed by fixation in 4% formaldehyde for 15 minutes. The sarcomeric structure of the adult heart was visualized with Alexa Fluor(tm) 568 Phalloidin (ThermoFisher Scientific).

### Mhc protein level quantification

For Mhc protein level quantification in tissues 1-week-old female mutant and control flies were stained under identical conditions as previously described (Alayari et al., 2009) using anti-Mhc (1:50, DSHB 3E8-3D3) and anti-mouse-Alexa488 (Jackson Labs, 1:500). Fly hearts were imaged using a Zeiss Imager M.1 microscope equipped with 25X dipping lens and identical imaging settings for each specimen (n=6-9 flies). Mean gray value (intensity) of 5 ROIs per tissue type (cardiomyocytes, ventral longitudinal muscle) were measured per fly using ImageJ and the average of mean fluorescent intensity per fly was calculated for each ROI.

### RNAscope

mRNA *in situ* hybridization for *CG14964* to count the number of transcripts per CM and body wall muscle and outlining of cells was done according to Blice-Baum *et al*. (Blice-Baum et al., 2018). The number of transcripts was counted in abdominal body wall muscles and CMs and normalized to the total area of each cell. Gapdh1/2 was used as a reference transcript (control). Similarly, we used a custom probe set to detect hMYOM2 (and *pericardin* as control probe) to show ectopic transcript expression of Mef2-Gal4-driven UAS-hMYOM2 in adult fly hearts.

### Sequence alignments and ortholog analysis

To compare *Drosophila* and Human protein sequences between CG14964 or bt with MYOM2, as well as bt or sls with TTN, we used the online LAST tool, which is based on a modified standard seed-and-extend approach and allows plotting amino-acid alignment in a dot plot manner (Kiełbasa et al., 2011).

### Statistical analysis

General statistical analyses were conducted using *R* or GraphPad Prism. Fly heart data presented as Cumming estimation plots were calculated using the *R* package *dabestr* estimating the unpaired mean differences to the control mean.

## Acknowledgments

We thank Sean Lal and Amy Li (Anatomy and Histology, School of Medical Sciences, Bosch Institute, University of Sydney, Australia) for providing control heart tissue, Ante Radocaj and Birgit Piep (Medical School of Hannover, Institute of Molecular and Cell Physiology, Hannover, Germany) as well as Sandra Schochardt-Schuster and Kerstin Mika (Cardiovascular Genetics, Charité - Universitätsmedizin Berlin, Berlin, Germany) for excellent technical assistance. We are deeply grateful to the patients and families for their cooperation.

## Author contributions

Conceptualization: E.A.-P., M.G., A.P., T.K., R.B., G.V., S.R.S.; Methodology: E.A.-P., T.N., G.V., T.K., R.B., G.V., S.R.S.; Software: M.G., G.V.; Validation: A.P., G.V.; Formal analysis: M.G., D.H., G.V.; Investigation: E.A.-P., T.N., M.G., O.O., A.P., D.H., F.M., C.M., G.V.; Resources: C.Ö., C.D.R., C.M., T.K., R.B., S.R.S.; Data curation: M.G., G.V.; Writing - original draft: E.A.-P., T.N., G.V., R.B.; Writing - review & editing: E.A.-P., T.N., M.G., A.P., T.K., R.B., G.V., S.R.S.; Visualization: M.G., O.O., A.P., D.H., F.M., C.M., G.V.; Supervision: T.K., R.B., G.V., S.R.S.; Project administration: T.K., G.V., R.B., S.R.S.; Funding acquisition: T.K., R.B., S.R.S.

